# High-quality proteins and RNAs extracted from exact same samples for proteomics and RNA-Seq analyses

**DOI:** 10.64898/2026.01.16.699903

**Authors:** Mathurin Fatou, Etienne Kornobis, Thibaut Douché, Karen Druart, Nicolas Puchot, Mariette Matondo, Marc Monot, Catherine Bourgouin

**Affiliations:** Institut Pasteur, Université Paris Cité, Biology of Host-Parasite Interactions, F-75015 Paris, France; Institut Pasteur, Université Paris Cité, Plate-forme Technologique Biomics, F-75015 Paris, France; Institut Pasteur, Université Paris Cité, Bioinformatics and Biostatistics Hub, F-75015 Paris, France; Institut Pasteur, Université Paris Cité, CNRS UAR 2024, Mass Spectrometry for Biology Unit, Proteomic Platform, F-75015 Paris, France

**Keywords:** Proteins and RNAs, omics analysis, exact same sample, mosquito

## Abstract

Back to the 1990’ the single step method developed by Chomczynski and Sacchi for RNA isolation was extended for sequential isolation of RNA, DNA and proteins from a same sample. Although the quality of the extracted RNA turned compatible with RNA-Seq analyses, the extraction of the protein fraction from the same sample was time-consuming and resulting in low yield and quality of proteins not compatible with LC-MS proteomic analyses. Here we report a novel procedure by isolating in parallel the protein fraction and the RNA fraction from the same exact minute mosquito samples. We provide evidence that each cognate fractions are compatible with LC-MS proteomic analysis on the one hand and RNA-Seq analysis on the other hand. This protocol is simple, time efficient and adequate for studies involving limited sample size and could be applied easily to a broad range of animal and human samples.

## Introduction

Advances in proteomics and transcriptomics technologies have greatly facilitated the comprehensive analyses of biological processes in various fields of biology. Combining Liquid Chromatography Mass Spectrometry (LC-MS)-based proteomics and RNA Sequencing (RNA-Seq) data has proven particularly powerful in identifying relevant pathways and differentially expressed genes in cells, tissues or whole organisms exposed to distinct conditions, notably in the context of host-pathogen interactions. In the case of cells in culture, protein and RNA extraction can be performed independently on replicated samples from the same passage. However, when working with limited tissue material, researchers often duplicate or split samples to separate extractions, raising concerns about the consistency of the starting material. To address this, some protocols perform sequential extraction from the same sample, typically isolating RNA first, followed by protein extraction using either commercial kits or custom lab methods (Chomczynski and Sacchi 1987; Chomczynski 1993; Hummon et al. 2007; Valledor et al. 2014). However, those protein extraction protocols are often time-consuming and may yield insufficient amounts for downstream proteomics analysis. Indeed, while the known TRI Reagent® or TRIzol^TM^ allow RNA extraction of good quality for RNA-Seq purpose, the vendors indicate that the extracted proteins are qualified for Western blots and some enzymatic assays (Molecular Research Center and Inc. 2017; Invitrogen and Thermo Fisher Scientific 2025).

In our quest for better understanding the response of different populations of *Anopheles gambiae* mosquitoes to the human malaria parasite, *Plasmodium falciparum*, we embarked in a comprehensive analysis of both proteins and mRNAs at play in *Plasmodium*-infected mosquito midguts by combining proteomic and transcriptomic analyses. Indeed, limited information exist on the relative stability (half-life) of proteins and mRNAs in *Anopheles* mosquitoes. It is usually accepted that at a defined time point during the interaction of *Plasmodium* with mosquito midgut cells, the detection of a regulated mRNA will match the presence-absence of the cognate protein. Conversely, looking at the proteome space may highlight stable proteins which corresponding mRNAs are no longer expressed at the studied time point. In addition, the correlation between mRNA and protein levels in organisms can vary according to several parameters as reviewed by Buccitelli and Selbach (Buccitelli and Selbach 2020). Notably, these authors highlighted that genome-wide mRNA and protein measurements can suffer from bias resulting from technical or biological replicates.

To limit such bias, we aimed at extracting RNA and proteins from the same exact mosquito midgut sample giving rise to molecules compatible for RNA-Seq and LC-MS proteomics analyses, meaning high quality and sufficient amount. Here, we present a streamlined and efficient protocol optimized for minute biological samples. The sample is first homogenized in urea, the gold standard denaturing solution in proteomics, under sonication assistance. The homogenate is then divided into equal parts for protein extraction on one side and RNA extraction on the other side. For the proteomic workflow, the sample undergo a standard bottom-up approach involving enzymatic digestion, peptide clean-up, and LC-MS analysis. RNA extraction from the urea homogenate is achieved through a simple protocol combining dilution in TRI Reagent® with a two steps purification on silica columns, yielding RNAs suitable for RNA-Seq applications. This protocol is simple, time efficient and particularly well-suited for studies involving limited sample size, enabling parallel proteomics and transcriptomics analyses from the same biological specimen.

## Results

### Overview of the technique

The objective of our protocol was to extract LC-MS compatible proteins and RNA-Seq compatible RNAs from the exact same sample of mosquito midguts. The diagram of the procedure is presented in Figure 1 and the detailed procedure in the M&M section.

**Fig. 1.**
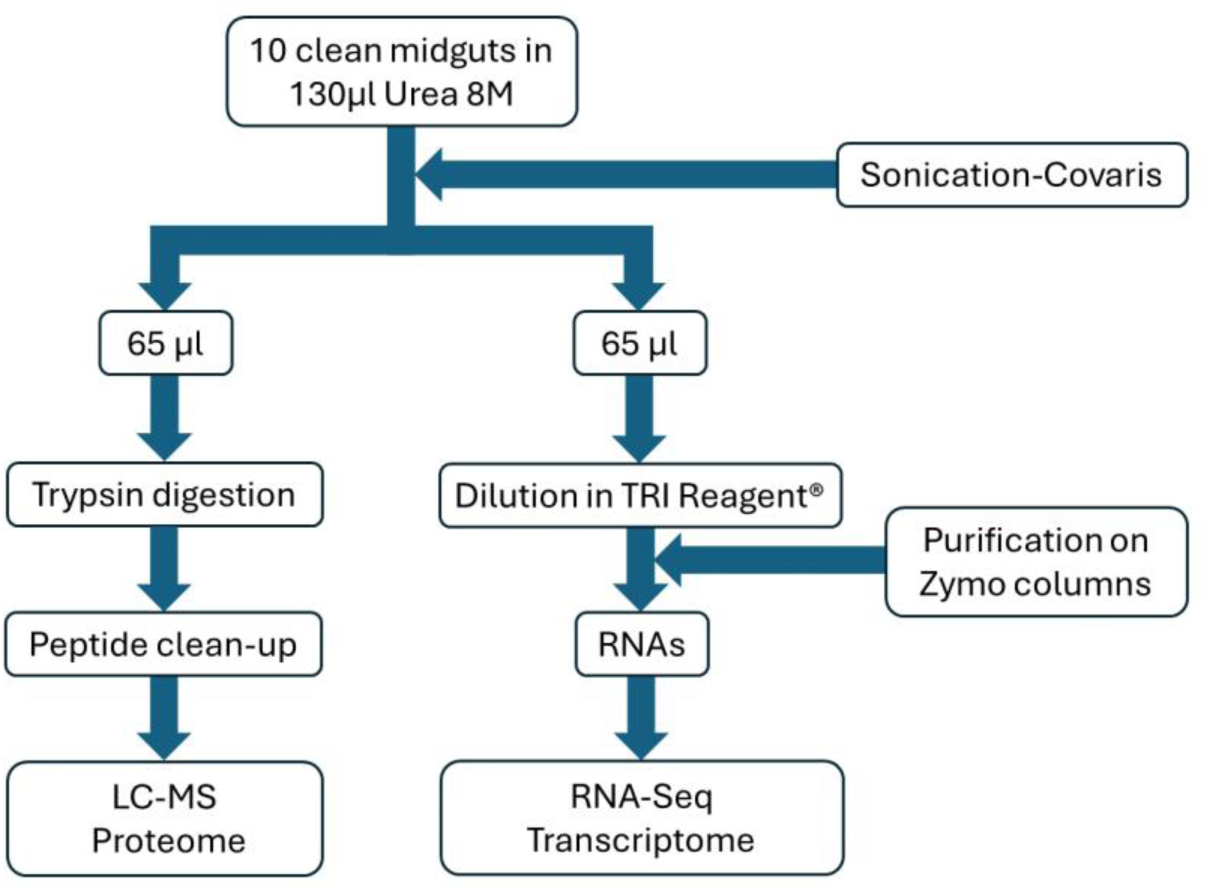
Protocol diagram. Ten mosquito midguts were pooled, homogenized by sonication and split to extract LC-MS compatible proteins and RNA-Seq compatible RNAs from the exact same sample.

The protocol was applied to four series of samples as described in Table S1. The first series corresponds to 6 pools of 10 midguts from uninfected mosquitoes, whereas the 3 other series correspond to 10 or 12 pools of 10 midguts from *P. falciparum* infected mosquitoes. Within a series, all samples were processed in parallel. Globally, the longest hands-on time part of the procedure is the step for RNA purification through silica columns as the result of the dilution of the initial urea extract in TRI Reagent®. Nevertheless, the length of this step remains very short close to only 2h for processing 10 samples (Table S2).

### Quality and quantity of the RNA samples for transcriptomic analysis

Half of each 10-midgut containing sample sonicated in urea were diluted in TRI Reagent®, and the RNAs were purified on Direct-zol columns (Zymo Research) and further cleaned using a dedicated kit from the same manufacturer. We found that the cleaning step was required to ensure high quality of total RNAs, as assessed using the Agilent 2100 Bioanalyzer. Indeed, the RIN (RNA Integrity Number) values exceed 6.0 for most samples (32 out of 38, Table 1), similar to the RIN values of RNA extracted from midguts sonicated in TRI Reagent® (Table S3). Further, the RNA profiles were very clear as presented in Fig. 2. Notably, the ribosomal RNA peaks were well preserved, keeping in mind that the Agilent RNA profile from insects deviates from the one of most eukaryotes (Winnebeck, Millar, and Warman 2010; Fabrick and Hull 2017; DeLeo et al. 2018; Ridgeway and Timm 2014).

**Fig. 2.**
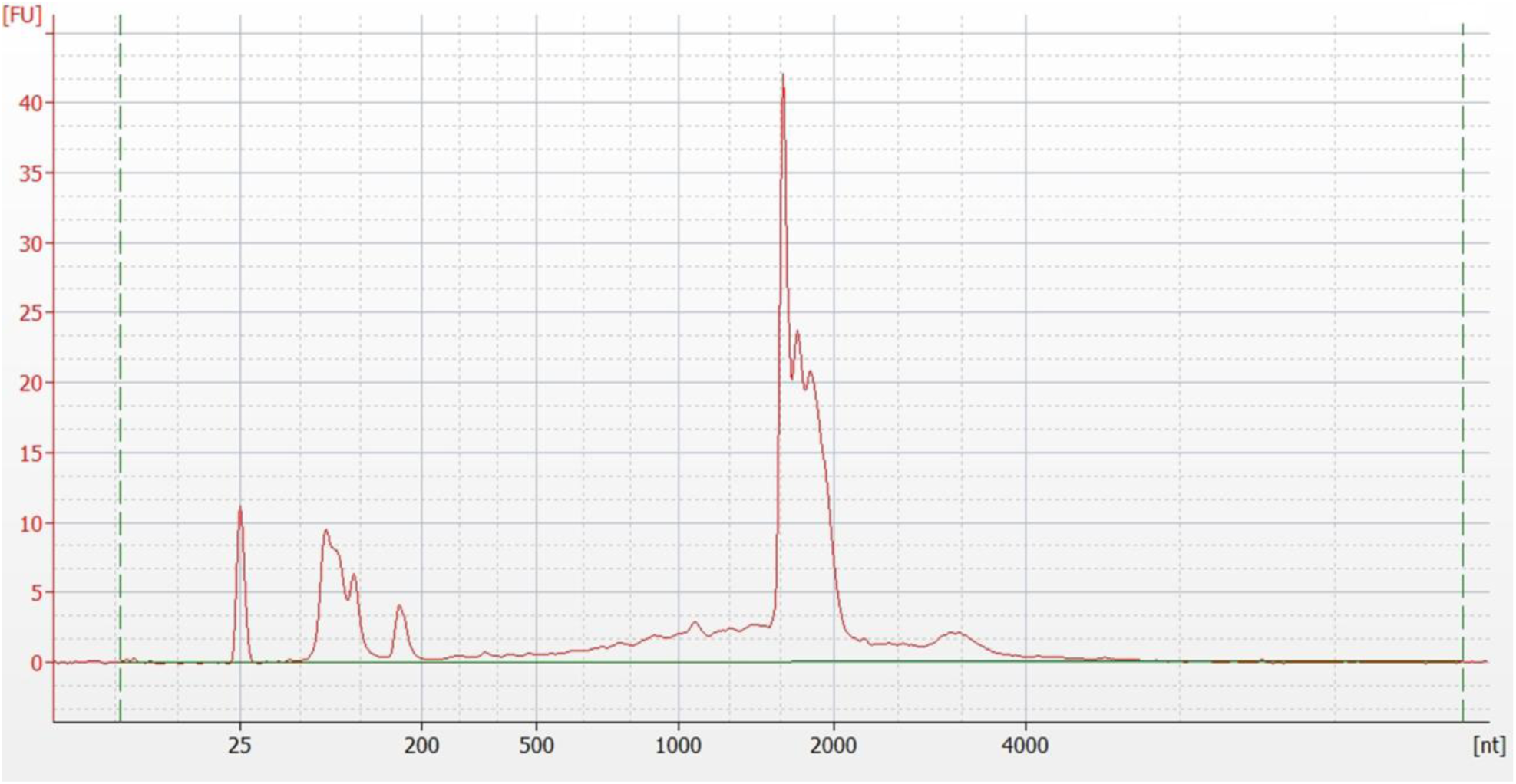
Electropherogram of sample U34 generated by 2100 Expert Agilent software. Marker can be seen at 25 nt, a combination of 18S and 28S RNA subunits at ±1900 nt, and a combination of 5S, 5.8S, and tRNAs at ±180 nt.

**Table 1.** Quantity and quality of extracted RNAs. The average total amount of RNAs was 1.7 µg (95% CI: 1.1–2.4 µg) in series 1, 0.6 µg (95% CI: 0.5–0.8 µg) in series 2, 0.7 µg (95% CI: 0.5–0.9 µg) in series 3 and of 0.7 µg (95% CI: 0.6–0.8 µg) in series 4. RIN: RNA Integrity Number.

| Series | Sample name | Total amount RNAs (µg) | RIN |
| --- | --- | --- | --- |
| 1 | U1 | 1.70 | 3.5 |
|  | U2 | 2.14 | 7.4 |
|  | U3 | 0.9 | 5.8 |
|  | U4 | 1.67 | 4.1 |
|  | U5 | 2.64 | 3.2 |
|  | U6 | 1.42 | 6.2 |
| 2 | U7 | 0.71 | 7.7 |
|  | U8 | 0.43 | 7.9 |
|  | U9 | 0.52 | 8.4 |
|  | U10 | 0.46 | 8.4 |
|  | U11 | 0.23 | 3.9 |
|  | U12 | 0.89 | 6.0 |
|  | U13 | 1.21 | 6.2 |
|  | U14 | 0.23 | 8.3 |
|  | U15 | 0.56 | 8.3 |
|  | U16 | 0.80 | 7.6 |
|  | U17 | 0.82 | 6.3 |
|  | U18 | 0.76 | 8.5 |
| 3 | U19 | 0.66 | 6.1 |
|  | U20 | 0.39 | 8.4 |
|  | U21 | 0.46 | 8.6 |
|  | U22 | 1.07 | 7.3 |
|  | U23 | 0.67 | 6.1 |
|  | U24 | 0.89 | 6.0 |
|  | U25 | 1.21 | 5.8 |
|  | U26 | 0.58 | 7.9 |
|  | U27 | 0.51 | 7.8 |
|  | U28 | 0.30 | 8.2 |
| 4 | U29 | 0.71 | 8.3 |
|  | U30 | 0.65 | 8.4 |
|  | U31 | 0.73 | 6.7 |
|  | U32 | 0.60 | 8.1 |
|  | U33 | 0.35 | 8.4 |
|  | U34 | 0.59 | 8.9 |
|  | U35 | 1.15 | 6.4 |
|  | U36 | 0.76 | 8.2 |
|  | U37 | 0.65 | 8.4 |
|  | U38 | 0.79 | 6.8 |

The amount of total RNAs extracted from an equivalent of 5 midguts varied from one sample to the other but reached values that are fully adequate to perform transcriptomic analyses. Indeed, our procedure produced an average of 1.7 µg of total RNAs in the first series, and between 0.6 µg to 0.7 µg for the 3 other series (Table 1).

Importantly, mRNA libraries using total RNAs extracted using our original procedure were suitable for downstream analysis according to the high alignment scores of a shallow sequencing presented in Fig. 3A and 3B, with low detection of ribosomal RNA (Fig. S1A and S1B).

**Fig. 3:**
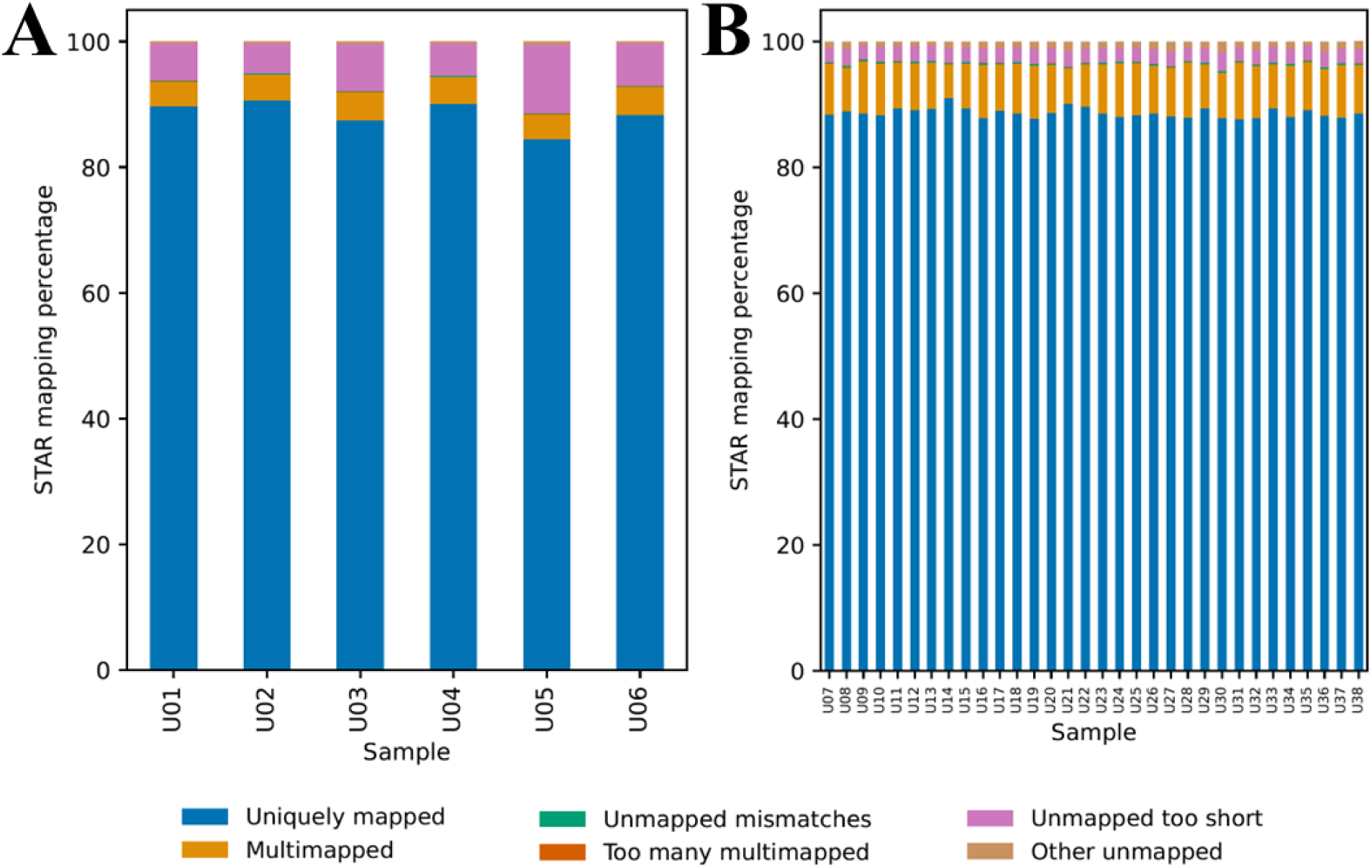
Percentage of mRNA alignments. The percentage of alignments was obtained using STAR 2.7.10a with the *Anopheles coluzzii* NCBI genome (GCF_943734685.1) as reference for a Iseq100 shallow sequencing runs (approximately 590k reads on average per sample). **A:** percentage of alignments from series 1. **B:** percentage of alignments from series 2 to 4.

### Quantity and quality of the protein samples for proteomic analysis

From the exact same samples, the cognate urea extracts (65µl) were processed for proteomic analysis using a standardized procedure for liquid samples under denaturing conditions. Half of the samples from the first series went to a single sonication using the softest parameters recommended for DNA shearing protocol provided on the Covaris company website. The second half of the samples went through an additional stronger sonication using the parameters recommended for protein extraction (supplementary text). The amount of extracted proteins and the number of identified proteins were similar among samples (Table S4, Fig. S2). Therefore, all the other samples (Series 2 to 4) went to a single soft sonication using the parameters described in the M&M section.

After LC-MS acquisitions and data treatment, the number of identified proteins was in the range of 4,300 proteins for series 1 and in the range of 6,100 for the 3 other series (Table 2). Although the first series used midguts from non-infected mosquitoes unlike the 3 other series that were from *Plasmodium*-infected mosquitoes, the difference in the number of identified proteins is most likely due to data acquisition using a more recent equipment rather than increased protein synthesis in the infected mosquito midguts.

**Table 2.** Quantity of extracted proteins. The number of identified proteins was obtained from an initial protein amount of 10µg. The 1.5 factor of identified proteins between series 1 and the 3 other series is due to the fact that two different generation of MS were used. Overall, from an equivalent of 5 midguts, the amount of extracted proteins was on average 22.9 µg (95% CI: 16.0–29.8 µg), 24.4 µg (95% CI: 19.3–29.6 µg), 22.9 µg (95% CI: 18.5–27.2 µg) and 23.6 µg (95% CI: 21.1–26.1 µg) in the first, second, third and fourth series, respectively.

| Series | Sample name | Protein amount<br>(µg) | Nb of identified<br>peptides | Nb identified<br>proteins |
| --- | --- | --- | --- | --- |
| 1 | U1 | 22.03 | 51,767 | 4,337 |
|  | U2 | 20.62 | 49,779 | 4,286 |
|  | U3 | 27.88 | 51,601 | 4,330 |
|  | U4 | 23.90 | 49,347 | 4,246 |
|  | U5 | 30.92 | 50,991 | 4,300 |
|  | U6 | 11.96 | 49,520 | 4,308 |
| 2 | U7 | 43.45 | 74,381 | 6,148 |
|  | U8 | 20.61 | 76,678 | 6,183 |
|  | U9 | 17.67 | 70,320 | 6,042 |
|  | U10 | 22.45 | 75,377 | 6,157 |
|  | U11 | 10.95 | 70,168 | 6,052 |
|  | U12 | 23.19 | 79,408 | 6,190 |
|  | U13 | 25.22 | 75,952 | 6,166 |
|  | U14 | 21.53 | 77,423 | 6,195 |
|  | U15 | 33.32 | 77,017 | 6,171 |
|  | U16 | 28.90 | 75,899 | 6,150 |
|  | U17 | 22.09 | 76,401 | 6,126 |
|  | U18 | 23.93 | 76,908 | 6,149 |
| 3 | U19 | 30.19 | 75,863 | 6,143 |
|  | U20 | 24.30 | 73,844 | 6,119 |
|  | U21 | 9.29 | 76,136 | 6,126 |
|  | U22 | 23.19 | 76,959 | 6,163 |
|  | U23 | 25.22 | 76,162 | 6,141 |
|  | U24 | 27.24 | 75,843 | 6,147 |
|  | U25 | 27.98 | 76,515 | 6,151 |
|  | U26 | 19.51 | 77,993 | 6,189 |
|  | U27 | 17.48 | 72,733 | 6,083 |
|  | U28 | 24.48 | 74,876 | 6,135 |
| 4 | U29 | 27.98 | 76,376 | 6,179 |
|  | U30 | 26.69 | 75,608 | 6,125 |
|  | U31 | 18.22 | 73,878 | 6,123 |
|  | U32 | 24.85 | 73,579 | 6,096 |
|  | U33 | 20.98 | 67,082 | 5,960 |
|  | U34 | 28.35 | 76,903 | 6,194 |
|  | U35 | 19.51 | 77,582 | 6,170 |
|  | U36 | 23.56 | 76,495 | 6,162 |
|  | U37 | 23.74 | 75,064 | 6,149 |
|  | U38 | 22.27 | 78,148 | 6,184 |

## Discussion

Here we described a novel procedure to obtain proteins and RNAs from the exact same sample in a simple and time-efficient manner that will be proteomics– and transcriptomics-compatible, respectively.

Based on an extensive literature search, nothing similar has ever been published. Indeed, we did a literature search using classical tools as PubMed, Scopus, Web of Science and generative AI tools (consensus and ChatGPT). Eighteen publications were obtained as outcome, but none were using our approach nor a similar one. Besides, to our knowledge no proteomics analysis complementing RNA-Seq data has ever been published in the *Plasmodium*-mosquito system. Our strategy is simple and efficient and overcomes the fact that acid-guanidinium-phenol based solutions are not suitable when protein profiling is the main aim of a study, mainly due to difficulties with the re-solubilization of the precipitated protein (Hummon et al. 2007).

Since the 2000s, RNA-Seq and LC-MS enable to reveal new parts of the genome that are transcribed and translated, to quantify RNAs and proteins as well as to detect transcripts and proteins with low expression (Woodland et al. 2025). Both transcriptome and proteome are rich in biological information and their crosstalk is overwhelmingly complex, hence, many studies combine RNA-Seq and LC-MS analyses to understand the interplay of the individual components of a biological system (Kumar et al. 2016).

Performing a multi-omics analysis from the exact same sample is obviously more robust than using different samples for each omic analysis as the phenotype between separate biological replicates might be substantially different. However, the challenge faced is the extraction of the desired proteins and RNAs, whereby there might be incompatibility for the subsequent workflows required to analyse them. Multiple strategies exist, but we came to the same conclusion than Woodland et al (Woodland et al. 2025) that the best possible approach for performing sample preparation for multi-omics analysis is to first homogenize the raw sample and then divide it equally into fractions before running each fraction through the intended downstream analysis. We even went further as RNA-Seq-compatible RNAs can be recovered from samples stored in urea 8M, the gold standard solution for protein extraction. If urea 8M had an effect at all on the extracted RNA, we didn’t observe it on the electropherograms assessing RNA integrity, and there was no difference with the electropherogram from a sample treated in TRI Reagent® only (not shown).

Recently, a study on the silkworm *Bombyx mori* used exact same samples for comparing both the transcriptome and the proteome across the moth developmental stages via RNA-Seq and LC-MS, respectively (Wilkens et al. 2025). In this study, each biological specimen was crushed in liquid nitrogen and dispatched for RNA or protein extraction. RNAs were extracted using a silica column based commercial kit similar to the one we used. However, proteins were processed using a combination of electrophoresis and in gel trypsin digestion for producing peptides for LC-MS analysis, similar to a previous study on *Caenorhabditis elegans* using exact same samples (Grün et al. 2014). This procedure is overall lengthier than the one we described here. Furthermore, the extraction yields of RNA and proteins were not mentioned making difficult to compare the efficiency of both procedures.

In conclusion, we developed a procedure with mosquito samples, and we deeply believe that our protocol can be applied to a broad range of animal or human samples: simplicity and time saving for collecting proteins and RNAs from the same exact starting material. It could be applied to cells in culture as well as any tissue including biopsies. It might be highly useful in medical science where the size of studied samples might be scarce as in cancer research. Furthermore, adaptation could be possible for semi-automation if large number of samples needs to be processed on the same day. An open question, that could be alleviated by testing, is the application of our procedure to plant samples where molecule extraction can be challenging due to the presence of polyphenols and polysaccharides.

## Material and methods

### Preparation of mosquito midgut samples

*An. gambiae* (Yaoundé strain) females were fed on non-infected human red blood cells (IcaReb, Institut Pasteur) or on *in vitro* produced *P. falciparum* gametocytes (CEPIA, Institut Pasteur) supplemented with human AB serum (Etablissement Français du Sang), using an artificial membrane feeding device (Hemotek Ltd, Blackburn, UK). Twenty-four hours later, mosquitoes were anesthetized by brief exposure to CO_2_. To remove external microorganisms, mosquitoes were then briefly submerged with 70% collected in a cell strainer and further washed in an additional 70% ethanol bath and twice with cold 1X PBS. From now on the mosquito samples were maintained on ice.

Blood containing midguts were isolated from each female by standard procedures using a stereomicroscope and transferred to cold PBS-containing wells of a 10-well slide placed on top of a cold sponge pad. The blood meal was removed using our specific procedure: the midgut was hold by its thin anterior part and dipped back and forth several times in the PBS solution, till the blood meal completely expels through the opening at the posterior part of the midgut and all remnants of the peritrophic matrix had disappeared (Fig. 4). Cleaned midguts from 10 mosquitoes were pooled into a protein low binding Eppendorf tube, containing 200µl cold PBS. After microcentrifugation (2,000 g, 10s), the midguts were suspended in 130 µl of urea 8M (Sigma-Aldrich, St-Louis, USA, ref U4883-25ML) supplemented with 50mM final concentration of protein-grade Tris (Sigma-Aldrich, St-Louis, USA, ref T2694-100ML) and stored at −80°C. According to the training of the experimenter, isolating 10 cleaned midguts free of any traces of blood meal and peritrophic matrix takes 30 to 40 min.

**Fig. 4.**
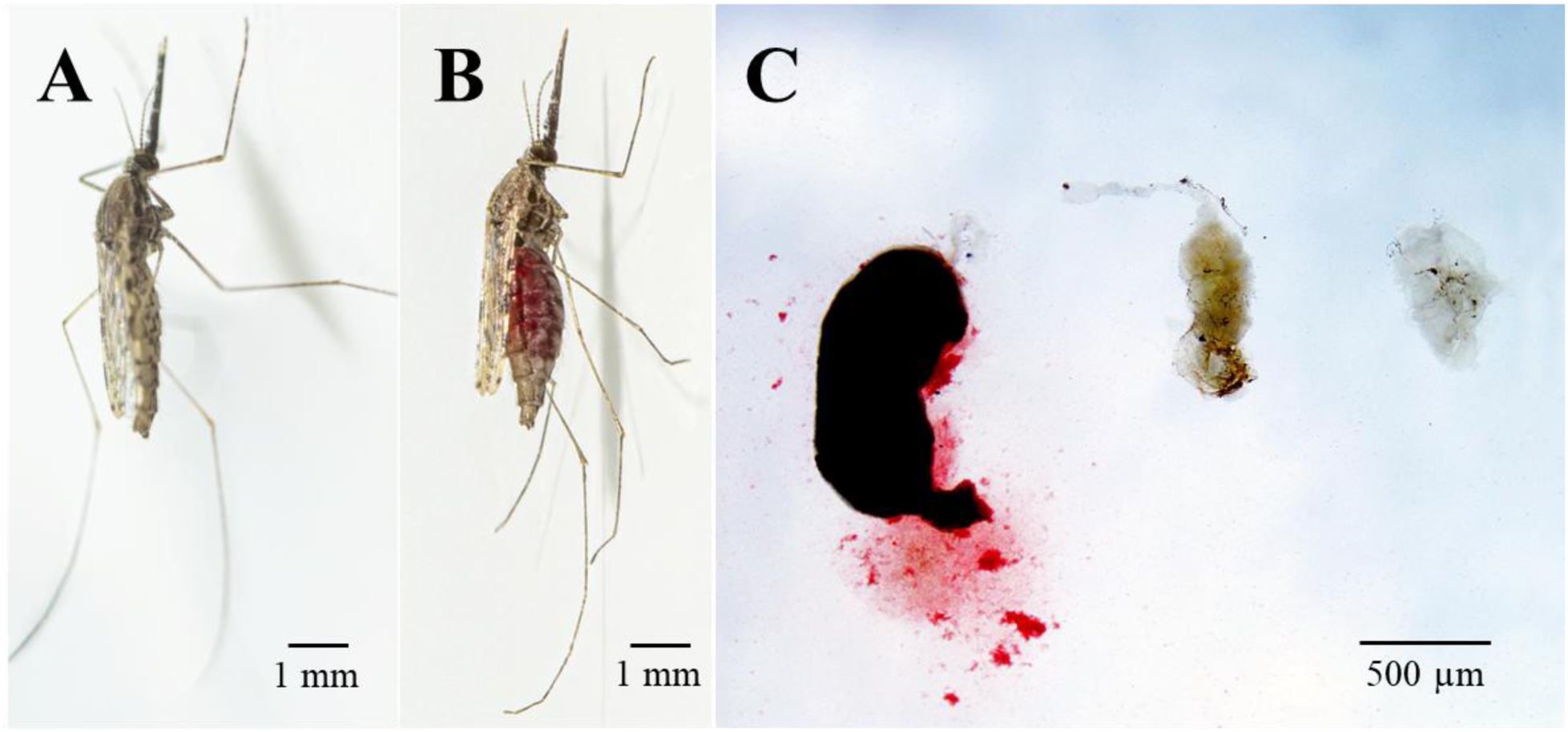
Dissected midgut from an *Anopheles gambiae* female. **A**: Unfed female. **B:** Blood-fed female. **C:** Status of the same midgut, from left to right: fully blood-filled, partially clean, clean. A and B pictures credit to François Gardy (Institut Pasteur).

### Sample sonication and protein extraction

Both proteins and RNAs were extracted from the exact same sample. Midguts suspended in 130 µl of 8M urea-50mM Tris-HCl pH 8.0, the gold standard solution for protein extraction, were gently thawed on ice, and the whole 130µl urea transferred into a microTUBE-130 AFA Fiber Screw-Cap 520216 (Covaris, Woburn, USA) under a hood. The sample underwent a sonication using a Covaris E220 (Covaris, Woburn, USA) using the following parameters: a peak power of 140 W, a duty factor of 2 %, 200 cycles per burst, a duration of 15s and a temperature of 5°C. After sonication, half of the sonicated content (i.e. 65 µl) was transferred to the proteomics platform of the Institute for Bradford dosage and MS analysis (see below). The other half was diluted 1:20 into TRI Reagent® (Sigma-Aldrich, St-Louis, USA, ref T9424-100ML), stored at −80°C to undergo RNA extraction on the following day.

### RNA extraction

After a gentle thaw on ice, each tube was centrifuged (Sigma-Aldrich, St-Louis, USA) to pellet the cellular debris (12000 g, 10 min), and each supernatant (1300 µl) transferred into a DNA low binding Eppendorf tube. Total RNA purification was performed using the Direct-zol^TM^ RNA Miniprep kit (Zymo Research, Irvine, USA, ref R2050) according to the manufacturer’s instructions. This kit is using silica columns for RNA purification. As the maximum volume a column could hold is 700 µl, 4 rounds of loading of 650 µl (1300 µl supernatant plus 1300 µl ethanol) and centrifugation (12,000 g, 1 min) were needed. A first DNAse treatment, was performed directly on the extraction column, as recommended by the manufacturer. Total RNA was eluted from the column twice with 25 μl of DNAse/RNAse free water. An additional DNAse treatment was performed on the final eluate, using the Invitrogen™ Ambion™ TURBO DNA-free kit (Thermo Fisher Scientific, Waltham, USA).

### RNA clean-up

RNA eluates (50 µl) were treated with the RNA Clean & Concentrator-5™ kit (Zymo Research, Irvine, USA, ref R1016) according to the manufacturer’s instructions and eluted in 10 μl of DNase/RNAse free water. Finally, 0.5 µl of RNaseOUT^TM^ Ribonuclease Inhibitor (Thermo Fisher Scientific, Waltham, USA) was added to the RNA samples that were kept on ice for the following step.

### RNA integrity and concentration

RNA integrity and concentration was assessed using the 2100 Bioanalyzer Instrument (Agilent, Santa Clara, USA) following the Agilent RNA 6000 Nano kit protocol without RNA heat denaturation. Electropherograms were generated after selecting the assay “Eukaryote Total RNA Nano Series II” on 2100 Expert software. RNA concentration and purity were further assessed using both Qubit assay and NanoDrop optical density measurement.

### RNA-Seq library preparation and Illumina sequencing

RNA preparation was used to build libraries using an Illumina Stranded mRNA library Preparation Kit (Illumina, USA) following the manufacturer’s protocol. Quality control was performed on an Agilent BioAnalyzer. The Illumina ISeq100 sequencer was used for a first test run of 6 pooled libraries (paired-end 150b reads, 1M reads per sample on average). The Illumina NextSeq2000 sequencer was used for the production run for 32 pooled libraries (single-end 65b reads, 52M reads per sample on average).

The RNA-seq analysis was performed with Sequana 0.18.2 (Cokelaer et al. 2017). We used the RNA-seq pipeline 0.20.0 (https://github.com/sequana/sequana_rnaseq) built on top of Snakemake 7.32.4 (Köster and Rahmann 2012). Briefly, reads were trimmed from adapters using Fastp 0.22.0 (Chen et al. 2018), then mapped to the *Anopheles coluzzi* (formerly named *An.gambiae* M) NCBI RefSeq genome (GCF_943734685.1) using STAR 2.7.10a (Dobin et al. 2013). Ribosomal content was evaluated by mapping reads to sequences annotated in the reference annotation as ‘rRNA’ using Bowtie2 (Langmead and Salzberg 2012). The featureCounts 2.0.1 program (Liao, Smyth, and Shi 2014) was used to produce the count matrix, assigning reads to features using corresponding annotation from NCBI with strand-specificity information. Quality control statistics were summarized using MultiQC 1.11 (Ewels et al. 2016). On average 42M reads for the production run were obtained and assigned to a gene, indicating sufficient material for future differential regulation analysis.

### Protein digestion

Sonicated samples were centrifuged at 16,000 g, 10 min, RT and protein assay (Bio-Rad, 500-0006) was performed using the standard procedure for microtiter plates of the user manual. For each sample, 10 µg of total proteins were homogenized in a final volume of 50 µL using 8M urea-50mM Tris-HCl, pH 8.0. Proteins samples were reduced with TCEP (Sigma – 646547) to a final concentration of 5 mM for 30 min and then alkylated with iodoacetamide (Sigma – I114) to a final concentration of 20 mM for 30 min in the dark. After a dilution by 9 of urea with 50 mM Tris-HCl, pH 8.0, a double digestion of the proteins were performed using Lys-C (Promega – V1671) with a ratio of 1:50 (enzyme:proteins) for 3H / 25°C and then using Sequencing Grade Modified Trypsin (Promega – V5111) with a 1:50 ratio (enzyme:proteins) for 16 hours / 37°C. Digestion was stopped with 1% formic acid (FA) final.

### Peptide clean-up

Digested peptides were desalted with the AssayMAP Bravo (Agilent Technologies) using 5μL bead volume C18 cartridge (Agilent – 5190-6532) and eluted with 80% acetonitrile (ACN), 0.1% formic acid (FA). Finally, the peptide solutions were speed-vac dried and resuspended with 2% ACN / 0.1% FA buffer. For each sample, absorbance at 280 nm was performed with a NanodropTM 2000 spectrophotometer (Thermo Scientific) to inject an equivalent OD=0.8 (series 1) on an Orbitrap Lumos tribrid or OD=0.175 (series 2 to 4) on an Orbitrap Astral.

### Mass spectrometry and database search

Two generation of mass spectrometer were used: an Obitrap lumos tribrid for samples U1 to U6 and a newly acquired Orbitrap Astral for the samples U7 to U38. For the samples U7 to U38, mass spectra were acquired in DIA mode with a nanochromatographic system (Vanquish Neo – Thermo Scientific) coupled online with an Orbitrap Astral mass spectrometer (Thermo Scientific). Peptides were injected into a 25cm C18 column (Aurora Ultimate TS, 75 μm ID, 1.7 μm beads – IonOpticks – HTS 9027205060) and eluted with a multi-step gradient from 6 to 18% solvent B in 18 min, 18 to 42% B in 4 min, 42 to 99% B in 4 min at a flow rate of 250 nL/min for up to 30 min. The composition for solvent A and B were 0.1% FA in water and 80% ACN, 0.1% FA respectively. Column temperature was set to 50°C.

MS data was acquired using the Xcalibur software with a scan range from 380 to 980 m/z with 240k Orbitrap resolution and 3ms maximum injection time for MS1. All DIA scans were set to 3ms maximum injection time with 2 Th windows. The normalized collision energy was set to 25 for HCD fragmentation. FAIMS CV was set to –48, with a total carrier gaz flow at 3.6 L.min^-1^. The features of the Obitrap lumos tribrid is described in the supplementary text.

Spectronaut 19 (Biognosys AG) was used for DIA-MS data analyses against the *Anopheles gambiae* Uniprot reference proteome (14,401 entries, download in 12/06/2024). The data extraction was performed using the default BGS Factory Settings. Briefly, for identification, both precursor and protein Qvalue were controlled at 1%. Quantification was performed at MS2 level and Qvalue was used for precursor filtering. Peptides were grouped based on stripped sequences. Imputation and cross run normalization was disabled.

## Acknowledgments

We wish to thank J-E Cluzel for mosquito production, GM Haustant from the Biomics Platform as well as France Génomique (ANR-10-INBS-09) and IBiSA (https://www.ibisa.net/). Acknowledgment goes as well to Sandrine Royer-Devaux (CeRIS-Institut Pasteur) for her help in literature searches. We also wish to thank Pr. Artur Scherf, head of the Biology of Host-Parasite Interactions unit, for his constant support.

## Author contributions

Conceptualization: CB, MMa, MMo

Methodology: CB, MMa, MMo, EK, TD

Investigation: MF, EK, TD, KD, NP

Visualization: MF, EK, TD, KD, NP, CB

Supervision: CB, MMa, MMo

Writing—original draft: MF, CB

Writing—review & editing: MF, CB, MMa, MMo, EK, TD, KD

## Data availability and materials sharing

All data needed to evaluate the conclusions in the paper are present in the paper and/or the Supplementary Materials.

Proteomic and transcriptomic data used in this paper can be provided upon reasonable request.

